# Camsap3-Mediated Microtubules Maintain Transzonal Projections Essential for Germline-Soma Communication during Ovarian Follicle Maturation in Mice

**DOI:** 10.1101/2025.09.26.678897

**Authors:** Akihiro Aikawa, Junya Ito, Mika Toya, Masamitsu Sato

## Abstract

Oocytes are surrounded and nursed by layers of maternal somatic granulosa cells (GCs) in ovarian follicles. Within a follicle, GCs extend actin-containing transzonal projections (TZPs) to an oocyte across the zona pellucida to establish communication. In contrast, microtubules have rarely been observed in some TZPs, and their significance in TZP organisation and functions for follicular maturation remain largely unknown. Here, using super-resolution microscopy, we visualised the presence of microtubules alongside F-actin in most TZPs. Knockout (KO) mice of the microtubule minus-end binding protein Camsap3 (calmodulin-regulated spectrin-associated protein 3) exhibited infertility with fewer developing late-stage follicles. In earlier stages of Camsap3-KO follicles, the number of TZPs was reduced compared to that in wild-type follicles, and microtubules in TZPs were disorganised, which led to a decrease in physical contact between GCs and oocytes. We also found that TZP morphology in wild-type follicles transforms over time during follicle development, and Camsap3 modulates TZP morphology by assisting the organisation of microtubules as well as F-actin. Our findings thus suggest that Camsap3-mediated microtubules play a crucial role in governing the number and morphology of TZPs to ensure successful follicle development for the production of fertile oocytes.

## RESULTS AND DISCUSSION

### *Camsap3*-KO female mice were infertile due to anovulation

*Camsap3* hypomorphic mutant mice *Camsap3^tm1a/tm1a^* reportedly displayed subfertility in both female and male mice, with a more pronounced effect in females^1^. Consistent with this, breeding records from the *Camsap3-dc* mutant line^2^ also indicated reduced fertility in females. To further clarify this phenotype, we employed *Camsap3*-null (*Camsap3*-KO) mice^3^. Female *Camsap3*-KO mice were completely infertile, producing no pups despite repeated mating with wild-type males, and showed no signs of pregnancy (Figure 1A). This infertility could not be attributed to disrupted oestrous cycles, as *Camsap3*-KO mice exhibited regular cycles every 4–5 days, comparable to that of wild-type controls.

**Figure 1.**
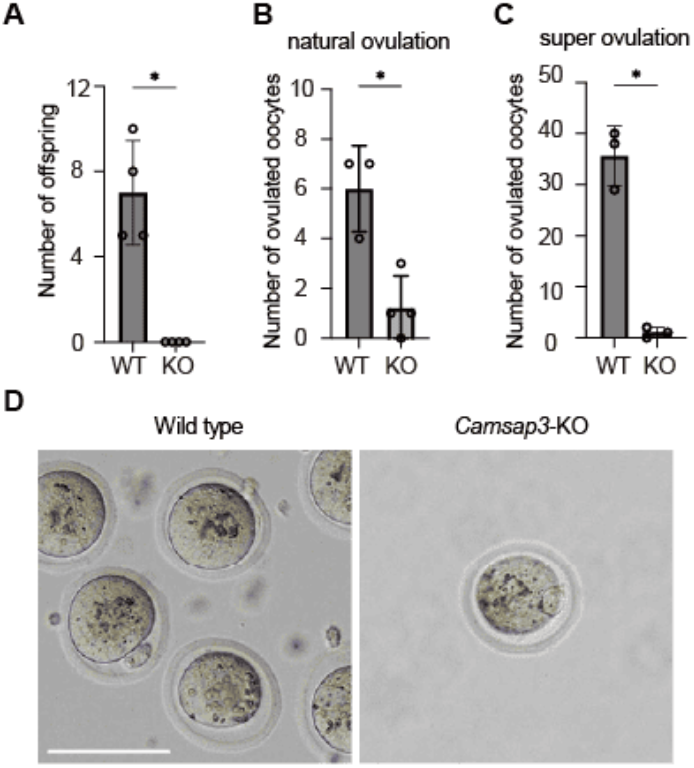
*Camsap3*-KO mice exhibited anovulation. (A) The number of offspring obtained by mating wild-type (WT) males with WT or *Camsap3*-KO females. (WT, n=4; KO, n=4). WT or *Camsap3*-KO females were housed with WT males for 30 days. *Camsap3*-KO females produced no pups, although vaginal plugs were observed, as in WT females. Bars, mean; error bars, s.d. *p < 0.05, two-tailed unpaired Student’s t-test. (B) The number of ovulated oocytes on the first day after mating (WT, n=3; KO, n=4). Bars, mean; error bars, s.d. *p < 0.05, two-tailed unpaired Student’s t-test. (C) Oocyte yield after superovulation (WT, n=3; KO, n=3). PMSG and hCG were administered 48 h apart; oocytes were collected 12 h after hCG. injection (D) Representative images of oocytes recovered from oviducts after superovulation. Scale bar, 100 μm.

Female mice are known to accept mating with males on the day of ovulation. To determine whether natural ovulation occurs in *Camsap3*-KO mice, we collected ovulated oocytes on the first day after mating, before implantation: 6.0 ± 1.4 oocytes were collected from wild-type mice, while 1.3 ± 1.1 oocytes from *Camsap3*-KO mice, suggesting that natural ovulation is impaired in the knockout mice (Figure 1B). These results suggest that infertility in *Camsap3*-KO female mice is due to a reduction in ovulated oocytes. To further assess ovulation defects, we induced superovulation with the hormones, pregnant mare serum gonadotropin (PMSG) and human chorionic gonadotropin (hCG), which promote follicle maturation and trigger ovulation. Upon superovulation, 36.7 ± 5.79 oocytes were recovered from wild-type mice, whereas 0.33 ± 0.47 from *Camsap3*-KO mice (Figure 1C), confirming that *Camsap3*-KO mice exhibit an anovulation phenotype, which is likely due to defects in follicle maturation in the ovaries. The few oocytes recovered from *Camsap3*-KO mice displayed morphology comparable to that of wild type, with a polar body and arrest at metaphase II accompanied by a meiotic spindle (Figure 1D, S1A).

### Delayed maturation after secondary follicles and increased atresia in *Camsap3*-KO mice

We examined ovarian follicle maturation in *Camsap3*-KO mice by analysing ovarian tissue sections. We classified follicles into five stages during maturation, namely primordial, primary, secondary, early antral, and Graafian follicles, based on their morphology^4^. After classification, we counted the number of follicles in each stage of the tissue sections from the whole ovaries.

At postnatal day (P) 4, all observed follicles were classified as primordial, with comparable numbers in wild-type and *Camsap3*-KO mice (4490 ± 700 in wild-type, 3335 ± 5 in *Camsap3*-KO; Figure 2A, 2B). Primordial follicles contain the Balbiani body (B-body), which is composed of the Golgi apparatus and is regulated by microtubules^5,6^. B-bodies play a crucial role in regulating the activation of primordial follicles^5,7^. As Camsap3 has been reported to localise to B-bodies^5^, we suspected its involvement in B-body organisation. However, the B-body morphology in *Camsap3*-KO was comparable to that in wild-type primordial follicles (Figure S2A), suggesting that there were no significant defects in primordial follicle formation.

**Figure 2.**
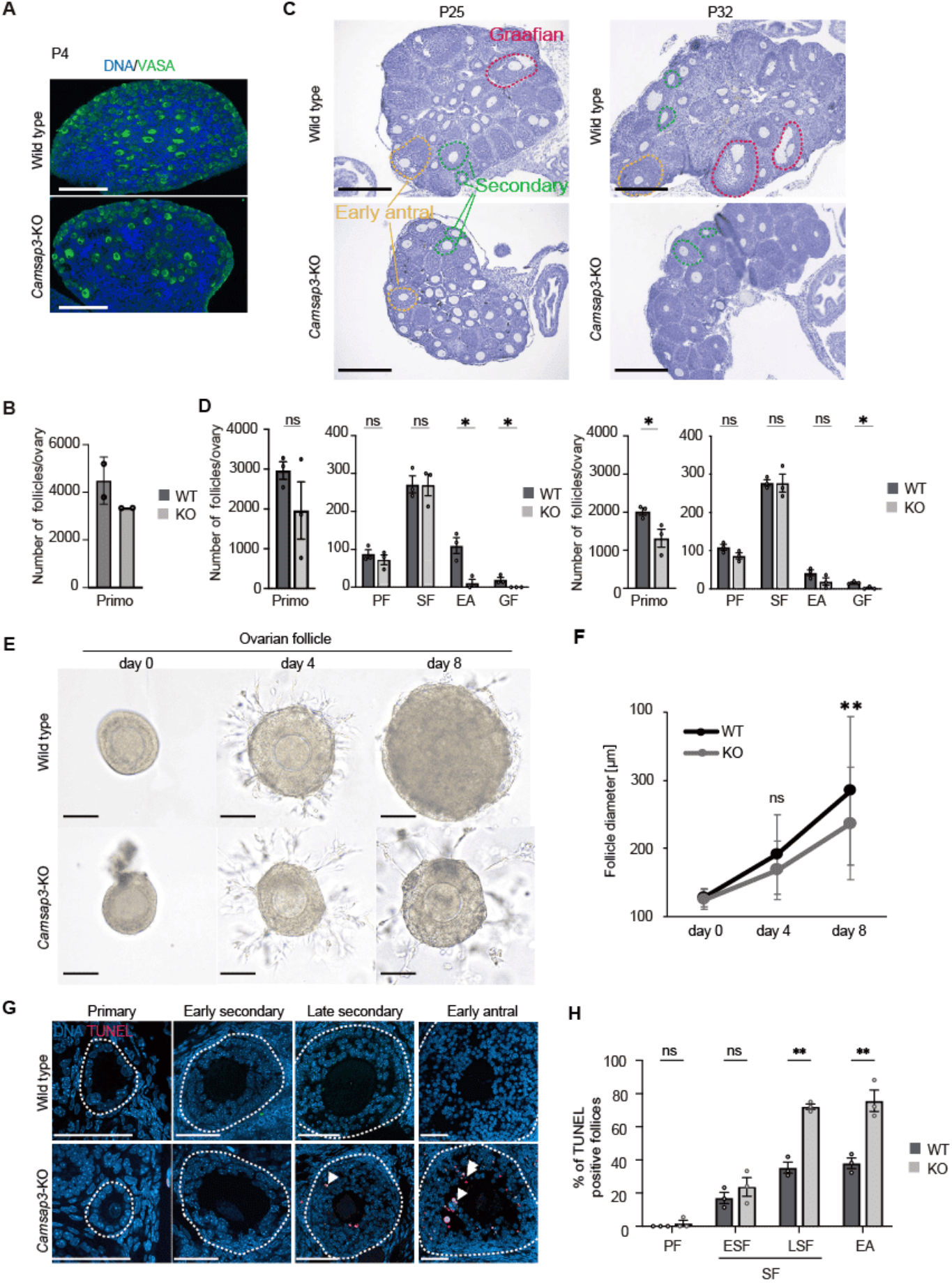
Delayed maturation after the secondary follicle stage and increased follicle atresia in *Camsap3*-KO mice. (A) Ovarian sections from P4 WT and *Camsap3*-KO mice stained for VASA and DNA. Scale bar, 50 μm. (B) The number of primordial follicles per ovary (WT, n=2; KO, n=2). Bars, mean; error bars, s.d. (C) Ovarian sections from P25 and P32 WT and *Camsap3*-KO mice stained with haematoxylin. Scale bar, 300 μm. (D) Follicle counts at each stage: primordial (Primo), primary (PF), secondary (SF), early antral (EA), and Graafian (GF) follicles (P25, P32: WT, n=3; KO, n=3). Bars, mean; error bars, s.d. *p < 0.05, two-tailed unpaired Student’s t-test. (E) Bright-field images of individual follicles encapsulated in Matrigel. Scale bar, 100 μm. (F) Average follicle diameter during Matrigel culture (WT, n=39 follicles; KO, n=40 follicles). Dots, mean; error bars, s.d. *p < 0.05, two-tailed unpaired Student’s t-test. (G)Sections of follicles at various developmental stages stained by TUNEL. Scale bar, 50 μm. (H) Percentage of follicles undergoing regression (TUNEL-positive GCs; white arrows) at the primary, early secondary, late secondary and early antral stages (WT, n=3; KO, n=3). Bars, mean; error bars, s.d. *p < 0.05, **p < 0.01, two-tailed unpaired Student’s t-test.

At P25, before the first oestrous cycle where all the stages of follicles are observed in wild-type ovaries^8^, the number of follicles in *Camsap3*-KO mice was comparable to that in wild-type mice up to the secondary follicle stage (271.3 ± 31.6 in wild-type, 229.6 ± 30.1 in *Camsap3*-KO; Figure 2C, 2D). However, follicles at more developed stages, the early antral and Graafian stages, were significantly reduced in *Camsap3*-KO mice (109.7 ± 29.6 in wild-type versus 11.3 ± 13.3 in *Camsap3*-KO in the early antral stage, and 20 ± 9.2 in wild-type versus 0 in *Camsap3*-KO at the graafian stage; Figure 2C, 2D). At P32, around the time of the first oestrous cycle, numbers of secondary follicles and early antral follicles in *Camsap*3-KO mice were comparable to those in wild-type mice (277.7 ± 11.6 in wild-type versus 276.3 ± 33.4 in *Camsap3*-KO at the secondary stage, and 41.7 ± 10.9 versus 18.7 ± 13.5 at the early antral stage; Figure 2C, 2D). However, Graafian follicles were significantly reduced in *Camsap3*-KO (16.0 ± 2.9 versus 2.3 ± 3.3; Figure 2C, 2D). It is possible that follicle development is delayed in *Camsap3*-KO mice, as their lighter body weight at P25 or P32^3^ potentially indicates reduced maturity compared with wild-type mice. To determine whether the developmental delay was due to slowed body growth rather than the loss of Camsap3 function in follicles, we examined the development of isolated secondary follicles *in vitro*. After eight days in culture, the secondary follicles isolated from wild-type developed to a diameter of 313.4 ± 115.3 μm, whereas those from *Camsap3*-KO mice showed limited growth to a diameter of 249.7 ± 82.7 μm, confirming that the developmental delay is due to the lack of the Camsap3 functions in follicles during the transition from secondary follicle to the early antral stage, rather than immature body conditions (Figure 2E, 2F).

At the age of 16 weeks, the period during which pregnancy is possible in mice, the proportion of follicles at various developmental stages in *Camsap3*-KO mice was comparable to that in wild-type follicles, although fewer Graafian follicles and corpora lutea were observed in *Camsap3*-KO mice (Figure S2B, S2C). Notably, no ovulated oocytes were observed at this stage in *Camsap3*-KO mice, unlike in wild-type mice, suggesting that follicle development in *Camsap3*-KO mice is impaired before reaching the Graafian stage. In ovaries, follicles that fail to develop undergo atresia, with GCs in these follicles undergoing apoptosis, as detected by TUNEL staining^9,10^. Therefore, we suspected that follicle regression was increased in *Camsap3*-KO mice. In P32 ovaries, the number of GCs undergoing apoptosis was significantly higher in late secondary and early antral follicles of *Camsap3*-KO mice (35.3 ± 4.8% in wild-type versus 72.2 ± 2.3% in *Camsap3*-KO in the late secondary stage, and 38.0 ± 4.5% versus 75.7 ± 9.1% in the early antral stage; Figure 2G, 2H). These results indicated that more follicles in *Camsap3*-KO mice regressed during development and rarely developed to the Graafian stage. Consistently, HE staining revealed an increase in follicles undergoing regression, characterised by nuclear condensation of GCs^11^ in *Camsap3*-KO mice (Figure S2D, S2E). The proportion of regressing follicles was significantly higher in *Camsap3*-KO mice than in wild-type mice, with 73.7 ± 12.6% versus 39.7 ± 1.5% in the late secondary stage, and 86.5 ± 9.7% versus 30.1 ± 15.1% in the early antral stage (Figure S2D, S2E). These results imply that the loss of Camsap3 decreases the developmental capability of GCs in the growing ovarian follicles.

### Camsap3 deficiency reduces transzonal projections between GCs and the oocyte

In ovarian follicles, layers of GCs surround an oocyte and extend cytoplasmic projections known as transzonal projections (TZPs), which penetrate the zona pellucida and reach the surface of the oocyte to establish direct contact between the GCs and the oocyte^12–14^. TZPs play a crucial role in supporting follicle maturation by facilitating intracellular communication and the transfer of essential molecules, including mRNAs and intracellular organelles such as mitochondria, from GCs to the oocyte^15–22^. TZPs are reportedly actin-rich filopodia-like structures, less than 5% of which also contain microtubules^14,23^. The function of microtubule-containing TZPs (tubulin-TZPs) and organisation of microtubules within them remain poorly understood.

In *Camsap3*-KO follicles, the number of TZPs detected by F-actin (actin-TZPs) was initially similar to that in wild-type follicles at the primary stage (5.8 ± 2.2 in wild-type vs. 6.1 ± 2.4 in *Camsap3*-KO), and then significantly reduced at the early secondary stage, which is right before the number of follicles begins to decline in *Camsap3*-KO (Figure 3A, 3B). This tendency lasted through the early secondary stage (5.8 ± 2.0 vs. 3.5 ± 2.4) until the early antral stage (4.8 ± 2.4 vs. 2.0 ± 1.2; Figure 3B and Figure S3A–S3D). In Graafian follicles, the number of actin-TZPs in *Camsap3*-KO was comparable to that in wild-type (Figure 3B). Given that the number of follicles per se declined in *Camsap3*-KO at the early antral stage, Graafian follicles counted here were rare ‘survivors’ (Figure 2C, 2D). As the number of actin-TZPs in those survivors was comparable to that in wild-type Graafian follicles, only follicles retaining actin-TZPs might be able to reach the Graafian stage. These observations suggest that Camsap3-mediated ‘microtubules’ play a more crucial role in maintaining TZPs than was previously understood.

**Figure 3.**
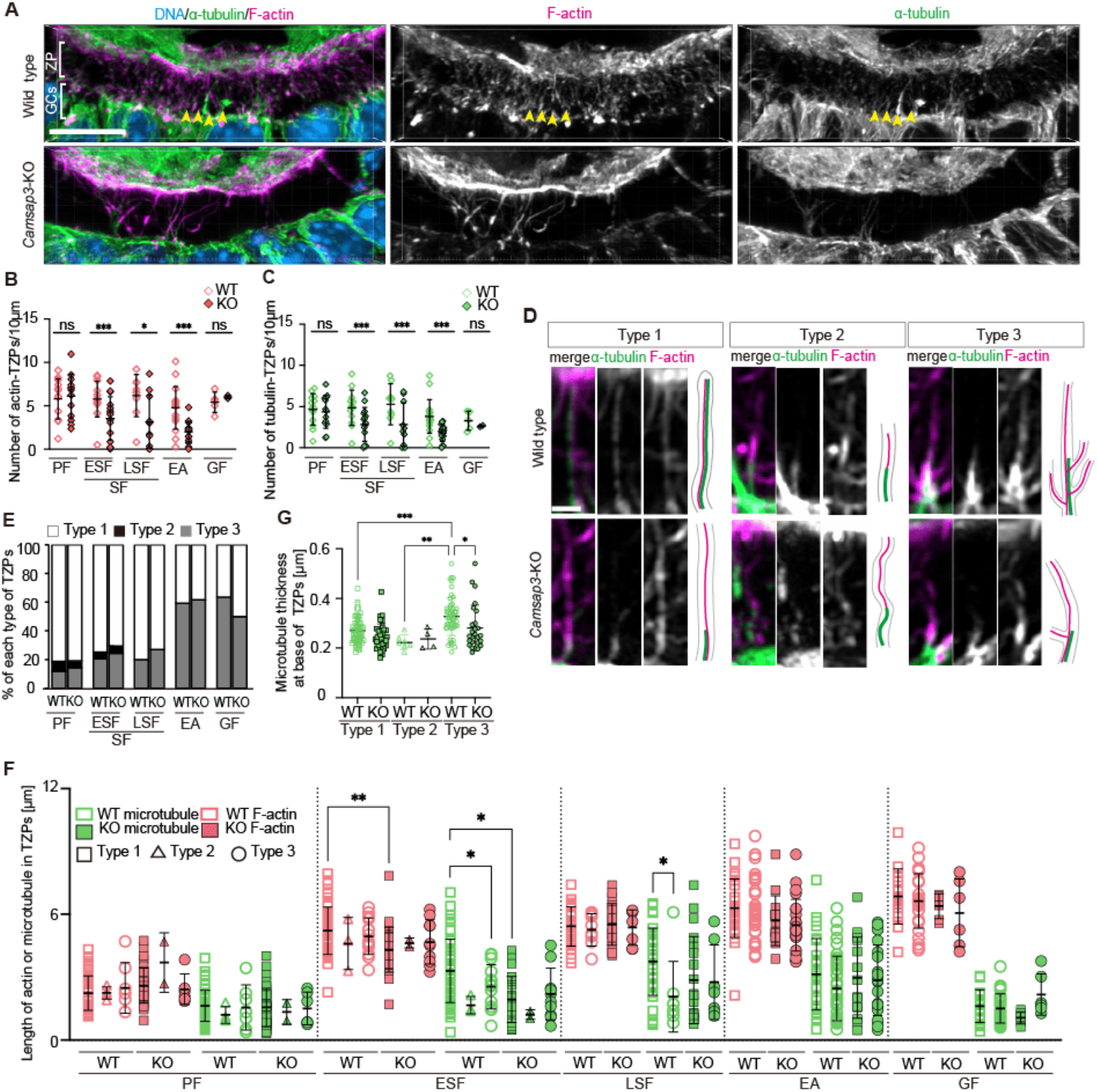
Most transzonal projections (TZPs) in wild-type follicles contain microtubules, and *Camsap3* deficiency reduces in the TZP number. (A) Three-dimensional immunofluorescence images of GCs and an oocyte at the early secondary stage, stained for α-tubulin, F-actin and DNA. Scale bar, 10 μm. (B–C) The average number of actin-TZPs and tubulin-TZPs per 10 μm at each follicle stage. The number of TZPs indicates the number of TZP roots formed from the apical surface of the cell. Dots, mean; error bars, s.d. *p < 0.05, **p < 0.01, two-tailed unpaired Student’s t-test. As shown in Fig. 2D, in *Camsap3*-KO mice at the early antral stage, Graafian follicles were rare ‘survivors’, and TZPs were assessed in these remaining follicles. (D) Representative images of three types of microtubule-actin arrangements. Scale bar, 0.5 μm. (E) The percentage of each TZP type (as defined in Figure 3D) by developmental stage. (F) F-actin and microtubule lengths of each TZP type at five stages of follicle development. Numbers of GCs examined: PF (WT, n =10 cells; KO, n = 10 cells), ESF (WT, n =10 cells; KO, n = 10 cells), LSF (WT, n =10 cells; KO, n = 10 cells), EA (WT, n =10 cells; KO, n = 10 cells), and GF (WT, n =10 cells; KO, n = 3 cells). Statistical analysis was performed using multi-group comparison of F-actin and microtubule datasets for each developmental stage. *p < 0.05, **p < 0.01 (G) Average microtubule thickness at the base of TZPs. Dots, mean; error bars, s.d. *p < 0.05, **p < 0.01, two-tailed unpaired Student’s t-test.

### Super-resolution inspection illuminates co-existence of microtubules and actin in most TZPs

In line with the significance indicated above, our super-resolution microscopy revealed that more than 80% of TZPs in wild-type follicles contained microtubules, in addition to F-actin (Figure 3A–3C). In wild-type follicles, the density of tubulin-TZPs peaked in the late secondary stage and then gradually decreased as follicle development progressed (Figure 3C).

In *Camsap3*-KO, the density of tubulin-TZPs was initially comparable to that in wild-type at the primary stage, but was lower than in wild-type from the early secondary to the early antral stages ––– the numbers of tubulin-TZPs per 10 μm of the luminal surface were: 4.6 ± 1.8 in wild-type vs. 4.3 ± 1.9 in *Camsap3*-KO at the primary stage; 4.9 ± 2.0 vs. 2.8 ± 2.0 at the early secondary stage; 5.3 ± 2.6 vs 2.8 ± 2.5 in late secondary stage; 3.8 ± 2.0 vs. 1.8 ± 1.0 at the early antral stage (Figure 3C). This suggests that Camsap3-mediated microtubules may play a role in TZPs in the secondary follicle stage, specifically or later. As the timing precedes the significant reduction of *Camsap3*-KO follicles in ovaries, defects in tubulin-TZPs caused by a loss of Camsap3 in secondary follicles impact subsequent follicle maturation in *Camsap3*-KO.

We characterised three distinct patterns regarding the internal architecture of TZPs that containing microtubules: type 1) microtubules and F-actin run alongside for most of the TZP length; type 2) microtubules are localised at the basal portion of the TZP, from the top of which F-actin extends towards the TZP tip; and type 3) F-actin branches off from the middle of a thick microtubule base like a ‘tree-trunk’ (Figure 3D). In both wild-type and *Camsap3*-KO follicles, all three types of TZPs were observed in the early stages of follicle development, with type 1 being the most dominant (over 70% of all tubulin-TZPs; Figure 3E). Type 2-TZPs diminished as follicle development progressed and disappeared by the late secondary stage, while the ratio of type 3-TZPs gradually increased up to over 60% at the Graafian stage (Figure 3E). The temporal transition of the ratios of the three TZP patterns during follicle development was equivalent in wild-type and *Camsap3*-KO follicles (Figure 3E).

The length of F-actin in TZPs increased as wild-type follicles grew (magenta, WT; Figure 3F), whereas the length of the microtubules decreased after the late secondary stage (green, WT), demonstrating that F-actin and microtubules were temporally rearranged according to stages to possibly fulfil specific functions for follicle maturation at each stage. In *Camsap3*-KO, microtubules, particularly those of type 1-TZPs, remained short in the early secondary follicles (green squares, Figure 3F). In wild-type follicles, microtubules at the bottom of type 3-TZPs were thicker than those in other types, whereas those in *Camsap3*-KO follicles were thin, similar to those in the other two types (Figure 3G). Thus, Camsap3 appears to play a role in the maintenance of microtubule length, particularly in type 1-TZPs during the early secondary stage, and in the thickness of microtubules in type 3-TZPs.

In type 2- and 3-TZPs, microtubules extend from GCs, with their distal ends connected to the proximal ends of the actin filaments. The actin filaments then extend from the tip or lateral surface of the microtubules towards the oocyte. A scheme including the microtubule-plus-end-binding protein CLIP-170 and the actin-nucleator formin may operate at the sites for the microtubule–actin interplay by analogy: CLIP-170 reportedly interacts with mDia1, a member of the formin family, and recruit it to the microtubule plus-end so that it accelerates actin polymerisation therefrom^24,25^. Under these conditions, CLIP-170 and mDia1 tracked the growing ends of the actin filaments^25^. Indeed, in the wild-type,13% of type-1, 4.0% of type-2, and 13% of type-3 TZPs displayed a CLIP-170 signal at the tip or lateral surface of actin filaments (Figure S3E–S3G), suggesting that microtubules in TZPs contribute to the nucleation and growth of actin filaments, allowing them to reach the oocyte more efficiently than TZPs consisting of microtubules only. Although CLIP-170 was detected in a limited manner at the microtubule-F-actin interface, this was due to transient localisation, as previously reported in vitro^25^.

Oocyte-secreted factors (OSFs), such as growth differentiation factor 9 (GDF9), regulate the formation of TZPs^14,26^. Oocyte-derived microvilli (Oo-Mvi) enrich and release OSFs to stimulate GCs^27^. In *Camsap3*-KO follicles, Oo-Mvi visualised with radixin staining were indistinguishable from those in wild-type, and GDF9 levels in the oocyte were comparable to wild-type in immunofluorescence (Figure S3H–S3J). These findings suggest that the lack of OSFs is unlikely to account for the reduction in TZPs observed in *Camsap3*-KO follicles. Rather, Camsap3 appears to contribute to the organisation and maintenance of TZP structure.

### TZPs contain Camsap3-associated microtubules with the mixed polarity that maintain GC–oocyte connection

Immunofluorescent staining showed that, in wild-type follicles, Camsap3 localised to TZPs and the apical region of GCs at all stages of follicle development (Figure 4A and Figure S4A–S4E). Camsap3 signals were also observed on the surface of the oocyte, although some appeared non-specific (Figure 4A). Camsap3 located at the base of microtubules near the apical surface of GCs, along the microtubules, and at the ends of microtubules towards the oocyte in all three types of TZPs (Figure 4B). Type 3-TZPs contained a significantly higher number of Camsap3 puncta at the bottom of the thick microtubule ‘tree-trunks’ than type 1- and type 2-TZPs (Figure 4B, 4C). Double-staining of Camsap3 which associates with the minus-end of non-centrosomal microtubules^28–30^, and the plus-end marker EB3 revealed that each TZP contained several microtubules of mixed polarity (Figure 4D). Magnified three-dimensional views from the apical side of GCs revealed that microtubules extended from the vicinity of intercellular junctions as well as from the middle of the apical surface in wild-type follicles (Figure S4F). In contrast, in *Camsap3*-KO GCs, TZPs tended to extend from the middle if present (Figure S4F). The remaining TZPs contained microtubules mediated by either the centrosome or Camsap2 but not Camsap1 (Figure S4G–S4L), suggesting that Camsap3, in cooperation with Camsap2 and the centrosome, stabilises the minus-ends of the microtubules to maintain their organisation.

**Figure 4.**
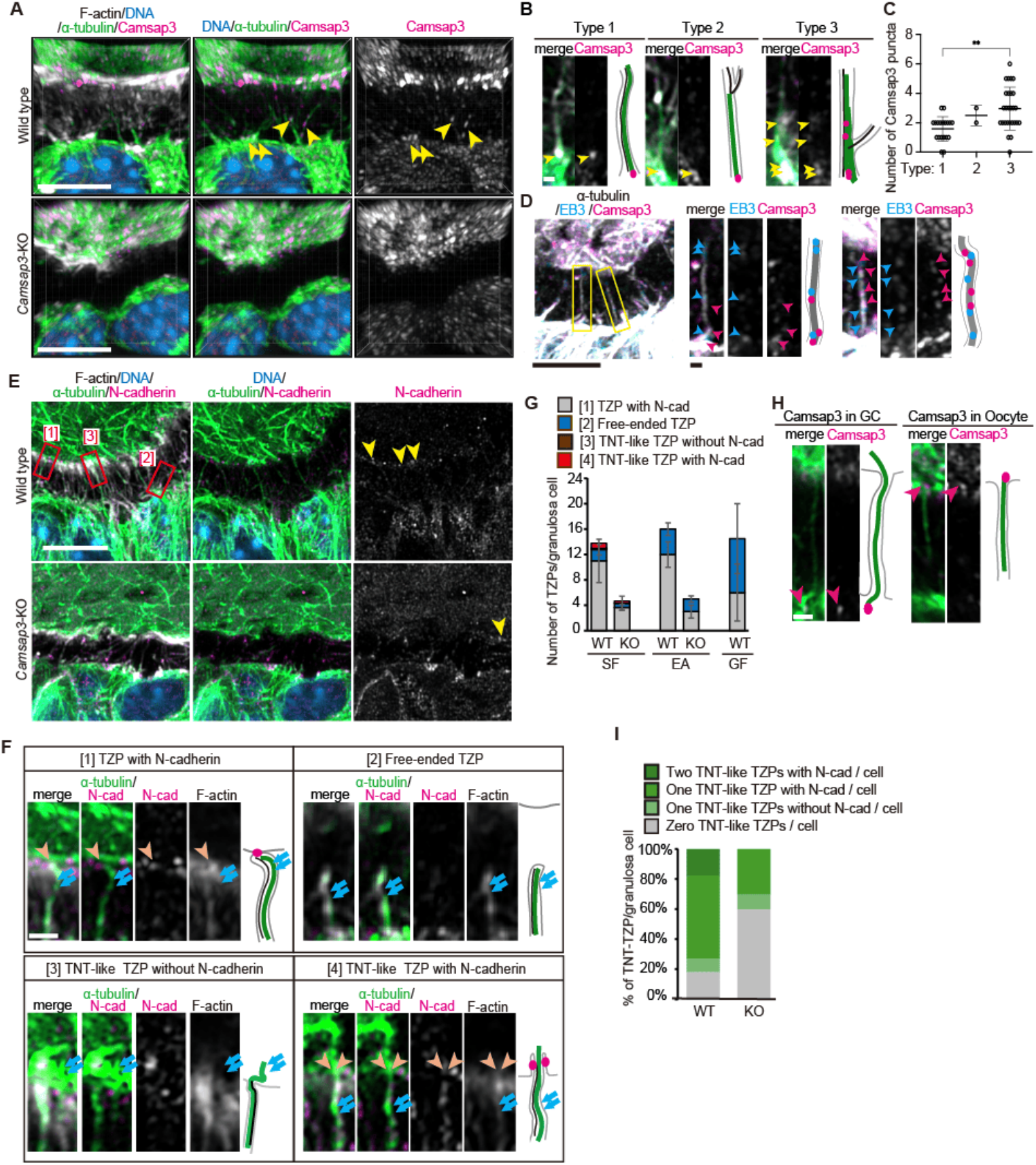
TZPs contain Camsap3-associated microtubules of mixed polarity that maintain oocyte-GC connections. (A) Three-dimensional Immunofluorescence images of GCs and an oocyte at the early secondary follicle stage, stained for α-tubulin and Camsap3. Scale bar, 5 μm. (B) Representative images of the three types of TZPs. Camsap3 localised to the base of the TZP and along the TZP. Scale bar, 0.5 μm. (C) The number of Camsap3 puncta localised per TZP for each type. (D) Immunostaining for Camsap3, EB3 and α-tubulin revealed that microtubules within TZPs displayed both plus and minus ends on the oocyte side. Scale bars, 5 μm, 0.5 μm. (E) Representative images of N-cadherin foci on the oocyte surface at the early secondary follicle stage. Scale bar, 5 μm. (F) Representative images of the contact area between an oocyte and TZPs: [1] N-cadherin localised to TZP tips in contact with the oocyte; [2] free-ended TZPs not connected to the oocyte; [3] TNT-like TZPs with microtubules penetrating the oocyte without N-cadherin; [4] TNT-like TZPs with microtubules penetrating the oocyte accompanied by N-cadherin. Scale bar, 0.5 μm. (G) Quantification of TZP types shown in (F) at each developmental stage. (H) Representative images of Camsap3 localisation to TNT-like TZPs. Camsap3 localised to the TZP base inside of GC or to the TZP tip inside the oocyte. Scale bar, 0.5 μm. (I) Proportion of TNT-like TZPs per GC.

At the TZP–oocyte interface, TZPs extended from GCs present N-cadherin, whereas the oocyte presents E-cadherin^31^, along with gap junctions displayed in both cell types^32^. Reflecting this, most TZPs of the secondary stage terminated at the oocyte surface, and some even penetrated the interior (Figure 4E, 4F), as previously described for Graafian follicles^33^. In the secondary stage, 74.1% of TZPs terminated at the oocyte surface, as indicated by the presence of N-cadherin ([1], Figure 4E–4G). In contrast, 7.3% of TZPs showed projection of the microtubule signal into the oocyte, with ([4]) or without ([3]) N-cadherin signals, suggesting that these TZPs may represent open-ended tunnelling nanotube (TNT)-like projections. TNT is a thin membranous structure that provides direct cytoplasmic connections between distant cells, enabling the transfer of vesicles and organelles^34^. TNT-like projections displayed Camsap3 at the end of penetrating microtubules (Figure 4H). Camsap3 was occasionally observed at the tip of TNT-like TZPs facing the oocyte, reflecting the mixed polarity of randomly oriented internal microtubules.

When the number of microtubules remained small in *Camsap3*-KO, GCs without any TNT-like projections substantially increased (60.0% in KO vs. 18.2% in WT; Figure 4I). Cell organelles such as mitochondria and mRNAs are known to translocate through TZPs^18–22^. We speculate that TNT-like TZPs may be selectively used for the translocation of these macromolecules because they cannot pass through cadherin-connected junctions. Thus, reduction of TZPs in *Camsap3*-KO, particularly the substantial loss of TNT-like projections, may cause severe deficiency in macromolecule translocation, which affects follicle maturation at an early stage.

This work demonstrates that Camsap3 contributes to follicle maturation through the organisation of TZPs, including TNT-like projections at the early stage of maturation, which is essential for GC–oocyte (soma– germline) communication. This explains the infertility phenotype caused by Camsap3 dysfunction in the female mice. *Camsap3*-KO follicles exhibited a reduced number of TZPs, and our super-resolution microscopy with optimised fixation for microtubules revealed that most TZPs contain microtubules. This defies the conventional view of TZP as actin-dominated specialised filopodia, since the existence of microtubules was underestimated by only ~5%. We thus unmasked the hidden significance of microtubules in TZP organisation. *Camsap3*-KO follicles in the secondary stage showed a substantial decrease in TNT-like TZPs accompanied by an increase in follicle atresia, suggesting that inefficient communication between GCs and the oocyte via tubulin-TZPs may primarily cause the symptoms, although additional contributions from other factors cannot be excluded. Although most *Camsap3*-KO follicles failed to reach the Graafian stage with a substantial loss of TNT-like TZPs, a few normal follicles with TZPs were found, in which Camsap2 may have compensated for the loss of Camsap3 function. Nonetheless, almost no ovulated oocytes were collected from *Camsap3*-KO mice, indicating that Camsap3 plays a crucial role in follicle maturation.

TZP contacts the oocyte surface via gap junctions and adherence junctions, through which molecules required for oocyte growth, such as pyruvic acid or cGMP, are transferred from GCs to oocytes^35,36^. Although it remains unclear how larger biomacromolecules, including mitochondria, are transported through cellular junctions, microtubule-prominent TNT-like TZPs may be key to their transport. Our observation of mitochondria in TZPs containing microtubules, but not in TZPs with actin filaments only, supports this hypothesis (Figure S4M). TNT-like tubulin-TZPs are observed only in the early stage of follicle development, suggesting that larger components essential for oocyte growth are efficiently transferred at this stage via microtubules that potentially facilitate directional transport.

This study demonstrated the temporal transition of TZPs during follicular development. In the early stages, the majority of TZPs were straight and contained both microtubules and F-actin (Figure S4N). GCs may use TNT-like TZPs to transfer macromolecules to oocytes. In contrast, in the later stages, branched TZPs with shorter microtubules increased. Branched TZPs reaching the oocyte surface may increase the area of physical contact through which relatively smaller molecules may be efficiently transferred, thereby controlling GCs to prepare for ovulation. The prevalence of thin and branched TZPs in the late stages may also facilitate timely degradation prior to ovulation. F-actin is the major component in the branches, but nonetheless ‘tree-trunk’ microtubules serve as the platform for the growth of the F-actin branches. Although TZPs thus transform over time, it is remarkable that Camsap3-mediated microtubules play a pivotal role in the assembly of both types of TZPs.

## Materials and Methods

### Mice

*Camsap3*-knockout (*Camsap3*-KO) mice used in this study were generated as described in our previous study (accession no. CDB0033E)^3^, in which the entire genomic region encoding Cmasap3 was deleted. N3 and N4 generation mice were used in this study. Backcrossing was performed using C57BL/6NCrSlc mice. Mice were housed in a specific pathogen-free (SPF) room. 8:00-20:00 lighted. The humidity was approximately 30-70 %, and the temperature was approximately 20°C. Use of these mice was approved by the Waseda University Animal Review Committee, and management and experiments were conducted in accordance with the guidelines and protocols provided by the committee.

### Genotyping

Genotyping of *Camsap3*-KO mice was performed by PCR as described previously^2^. Tail biopsies were used to extract template DNA. To distinguish between wild-type and *Camsap3*-KO alleles, three primers (P1, P2, P3) reported in Mitshuhata et al., 2021^3^ were used. PCR amplification with P1/P2 generated a 411-bp product corresponding to wild-type allele, whereas amplification with P1/P3 yielded a 614-bp product corresponding to the *Camsap3*-KO allele. The P1/P3 region in wild-type mice was not amplified under the PCR conditions used.

### Fertility test

Female mice were tested between 8 and 17 weeks of age, and male mice were tested from 8 weeks of age onwards. For crossbreeding, males larger than females were selected. One female and one male were co-housed for one month, and females were examined daily for the presence of vaginal plugs. The mating combinations were: (1) ♂ wild-type × ♀ wild-type, (2) ♂ wild-type × ♀ *Camsap3*-KO. Successful copulation was confirmed by the presence of a vaginal plug, after which the pair was separated. Mice typically give birth 19 days after successful copulation.

### Natural ovulation and Superovulation

For superovulation treatment, 8-weeks-old wild-type and *Camsap3*-KO mice were injected with 10 IU PMSG (ASKA Pharmaceutical) at 17:00, followed 48 h later by 5 IU hCG (ASKA Pharmaceutical). Mice were dissected 12 h after hCG administration. Cumulus cells were removed using 50 mM hyaluronidase (Sigma), and the number of oocytes were counted. For natural ovulation, oocytes were collected the morning after confirmation of a vaginal plug.

### *In vitro* growth of ovarian follicle

As previously reported^27^, follicles were grown for maturation in a three-dimensional culture system using Matrigel (Becton Dickinson). Matrigel was mixed with culture medium (3:1) on ice; the medium consisted of FSH (10 mIU/ml; Merck), ITS (Sigma), and 10 % FBS (NICHIREI) in MEMα supplemented with GlutaMax (Thermo Fisher). Drops of the mixture (20 μl) were placed at the bottom of a 35-mm Petri dishes and incubated for 20 min to solidify. Follicles of approximately 130 μm in diameter were isolated from 3-week-old mice in MEMα and embedded in the gel using a mouth pipette. Cavities generated during the process were sealed with 2 μl of additional Matrigel. Each dish was filled with 1 ml of the culture medium and incubated at 37°C with 5 % CO_2_ for 8 days, with half of the medium replaced every other day.

### Haematoxylin staining

Tissue samples were prepared as previously described^37^, with minor modifications. Glass slides with paraffin sections were first soaked in xylene for 5 min, followed by soaked twice for 3 min each. The samples were sequentially immersed twice in 100 % ethanol for 3 min, 90 % ethanol for 3 min, 80 % ethanol for 3 min, followed by 70 % ethanol for 3 min. After immersion in haematoxylin solution (Wako) for 3 min, the sections were washed in tap water for a few seconds and then transferred to tap water at 30°C for 15 min. The samples were washed with 100 % ethanol. The samples were then soaked in ddH_2_O with manual shaking constantly for 15 min and mounted with glycerol.

### TUNEL assay

Paraffin sections of 5 μm thickness were used. For deparaffinisation, the sections on glass slides were placed on a heating plate at 55°C for 30 min, followed by two sequential 15-min immersions in xylene. To remove xylene, the samples were sequentially immersed twice in 100 % ethanol for 5 min, 90 % ethanol for 5 min, 80 % ethanol for 5 min, and 70 % ethanol for 5 min. The samples were then immersed in ddH_2_O for at least 20 min. The samples were treated with Proteinase K solution (10–20 μg/ml in 10 mM Tris/HCl, pH 8) and incubated at 37°C for 30 min, followed by wash with PBS for three times. The samples were dried up, and 50 μl of TUNEL reaction solution (Merck) was applied. The samples were incubated at 37°C for 60 min in the dark. VECTASHIELD mounting medium (Vector laboratories) was used for the encapsulation.

### Immunofluorescent staining

Tissue samples were prepared as previously described^2^, with minor modifications. Ovarian tissue was fixed by immersion in 2 % PFA/50 mM Sorbitol/PEM for 1 h at room temperature and then washed three times for 10 min each in PEM buffer. For cryoprotection, tissues were sequentially transferred to sucrose/PEM solutions: 15% for 2 h at 4°C, 20 % overnight at 4°C, and 30 % for 5 h at 4°C. Samples were embedded in OCT compound (Sakura Finetek Japan) and snap-frozen, in liquid nitrogen, then stored at –80°C. Frozen blocks were sectioned at 5 μm using a CM1950 (Leica).

Glass slides with frozen ovarian sections were incubated in 0.1 % Triton X-100/PEM for 10 min at room temperature, followed by permeabilisation in 0.2 % Triton X-100/PEM for 10 min. Samples were then washed in 0.1 % Triton X-100/PEM with shaking for 10 min. Blocking was performed with 3 % bovine serum albumin (BSA)/0.1 % Triton X-100/PEM for 1 h at room temperature. Primary antibodies, diluted in the sane buffer, were applied overnight at 4°C. After incubation, samples were washed three times for 15 min each in 0.1 % Triton X-100/PEM. Secondary antibodies, diluted in 3 % BSA/0.1 % Triton X-100/PEM, were applied for 2 h at room temperature, followed by sequential washes in 0.1 % Triton X-100/PEM for 5, 10, and 15 min. Samples were mounted in ProLong Glass (Thermo Fisher) or the VECTASHIELD mounting medium.

Oocytes were washed in 0.2 % NGS/PEM for 2 min three times, then fixed in 2 % PFA/0.05 M sorbitol/PEM for 1 h. After fixation, samples were washed again in 0.2 % NGS/PEM for 2 min three times, followed by permeabilisation in 0.25 % Triton X-100/PEM for 15 min. Oocytes were then washed three times in drops of 0.1 % NGS/0.01% Triton X-100/PEM and incubated in a fresh drop of the same solution for 10 min three times. Blocking was performed in 2 % NGS/0.1 % Triton X-100/PEM for 1 h. Antibodies were diluted in the same solution and then applied for 2 h at room temperature. Samples were washed sequentially in 0.1 % Triton X-100/PEM for 5, 10, and 15 min at room temperature and mounted in the VECTASHIELD mounting medium.

### Microscopy

Images were acquired with following microscopes. BZ-X710 All-in-One Fluorescence Microscope (Keyence) equipped with the objective lenses Nikon Plan Apo λ 2×, 10×, 20×, and Nikon CFI Flour 4×. Images were then processed using the BZ X Analyzer software (version 1.3.1.1). LSM980 laser scanning confocal microscope with Airyscan 2 (Zeiss), equipped with the objective lens Zeiss Plan apo 63×. Z-slices were obtained using a 0.15 μm step. Images were processed using the software ZEN (version 3.4.91.00000). Nikon ECLIPSE Ti2 (Nikon) equipped with the objective lenses Nikon plan Flour 10×, 20× and 40× was used for DIC images of oocytes.

### Image Analysis

Fiji (version 1.53)^38^, Imaris (version 10.1.1) and ZEN (version 3.4.91.00000) were used for image analyses. Imaris was used to analyse the TZP length, microtubule thickness, and Camsap3 localisation.

### Follicle count

Sections of the entire ovary, sliced at 5 μm each, were stained with haematoxylin, and microscopic images were acquired. The numbers of primary, secondary, early antral, and Graafian follicles were counted, and the number of follicles per ovary was calculated. Primary follicles were defined as those contain a single layer of GCs, secondary follicles contain two or more layers of GCs with no visible follicular cavities, early antral follicles contain three or more layers of GCs with small follicular cavities, and Graafian follicles contain three or more layers of GCs with a single, united follicular cavity^27^.

The number of primordial follicles was determined as follows: the entire ovary was sliced into 5-μm slices, and every five section was selected for counting. The primordial follicles in the whole section were counted, and the number was multiplied by five, to estimate the number of primordial follicles per ovary^27^.

### TZP count

Ovarian sections containing each developmental stage of follicles were stained for α-tubulin and F-actin for imaging of TZPs. In the images, a line was drawn above the apical surface of the GCs in parallel. A plot profile of Zen was applied for drawing the line, and the number of peaks of the fluorescence intensities of F-actin and α-tubulin were measured. The ratio of the number of peaks to the total line length was calculated.

### GDF9 intensity

Secondary and early antral follicles were included in the analyses. Average fluorescence intensities of GDF9 inside an oocyte and in the zona pellucida were both measured. Then, the relative fluorescence intensity of GDF9 was calculated by subtraction of the value in zona pellucida from that in the oocyte.

## Statistical analysis

All the experiments were performed at least twice. Data were analysed using Student’s t-test, or one-way ANOVA with Prism software (version 9.4.1 (458); GraphPad).

## Acknowledgments

We thank Masatoshi Takeichi (RIKEN BDR) for experimental materials, valuable discussions, and helpful suggestions. We are also grateful to Erina Kuranaga for constructive comments on the manuscript, and to Kanako Tsuzuki and Sonoko Saji for their contribution to the initial stages of this study. A.A. was supported by JST SPRING, Grant Number B2R101263201. This work was supported by JSPS KAKENHI Grant Numbers 25K09625 to M.T., and 16H04787, 18K19347, 23K27173 and 25H02582 to M.S. Support also came from Daiichi Sankyo Foundation of Life Science to M.S, Ohsumi Frontier Science Foundation to M.T., and Waseda University grants for Special Research Projects Grant Numbers 2022C-170, 2023Q-012, 2024R-029 (to M.T.), 2017B-243, 2020R-038, 2023C-167, 2024C-490, 2025R-070 and 2025C-154 (to M.S.).

## References

1. Robinson, A.M., Takahashi, S., Brotslaw, E.J., Ahmad, A., Ferrer, E., Procissi, D., Richter, C.-P., Cheatham, M.A., Mitchell, B.J., and Zheng, J. (2020). CAMSAP3 facilitates basal body polarity and the formation of the central pair of microtubules in motile cilia. P Natl Acad Sci Usa 117, 13571–13579. 10.1073/pnas.1907335117.

2. Toya, M., Kobayashi, S., Kawasaki, M., Shioi, G., Kaneko, M., Ishiuchi, T., Misaki, K., Meng, W., and Takeichi, M. (2016). CAMSAP3 orients the apical-to-basal polarity of microtubule arrays in epithelial cells. Proc National Acad Sci 113, 332–337. 10.1073/pnas.1520638113.

3. Mitsuhata, Y., Abe, T., Misaki, K., Nakajima, Y., Kiriya, K., Kawasaki, M., Kiyonari, H., Takeichi, M., Toya, M., and Sato, M. (2021). Cyst formation in proximal renal tubules caused by dysfunction of the microtubule minus-end regulator CAMSAP3. Sci Rep-uk 11, 5857. 10.1038/s41598-021-85416-x.

4. Pedersen, T., and Peters, H. (1968). PROPOSAL FOR A CLASSIFICATION OF OOCYTES AND FOLLICLES IN THE MOUSE OVARY. Reproduction 17, 555–557. 10.1530/jrf.0.0170555.

5. Lei, L., Ikami, K., Miranda, E.A.D., Ko, S., Wilson, F., Abbott, H., Pandoy, R., and Jin, S. (2024). The mouse Balbiani body regulates primary oocyte quiescence via RNA storage. Commun. Biol. 7, 1247. 10.1038/s42003-024-06900-4.

6. Niu, W., and Spradling, A.C. (2022). Mouse oocytes develop in cysts with the help of nurse cells. Cell. 10.1016/j.cell.2022.05.001.

7. Lei, L., and Spradling, A.C. (2016). Mouse oocytes differentiate through organelle enrichment from sister cyst germ cells. Science 352, 95–99. 10.1126/science.aad2156.

8. Zheng, W., Zhang, H., Gorre, N., Risal, S., Shen, Y., and Liu, K. (2014). Two classes of ovarian primordial follicles exhibit distinct developmental dynamics and physiological functions. Hum Mol Genet 23, 920–928. 10.1093/hmg/ddt486.

9. Li, J., Kim, J.-M., Liston, P., Li, M., Miyazaki, T., Mackenzie, A.E., Korneluk, R.G., and Tsang, B.K. (1998). Expression of Inhibitor of Apoptosis Proteins (IAPs) in Rat Granulosa Cells during Ovarian Follicular Development and Atresia 1. Endocrinology 139, 1321–1328. 10.1210/endo.139.3.5850.

10. Kim, J.-M., Boone, D.L., Auyeung, A., and Tsang, B.K. (1998). Granulosa Cell Apoptosis Induced at the Penultimate Stage of Follicular Development is Associated with Increased Levels of Fas and Fas Ligand in the Rat Ovary1. Biol. Reprod. 58, 1170–1176. 10.1095/biolreprod58.5.1170.

11. Hirshfield, A.N. (1988). Size-Frequency Analysis of Atresia in Cycling Rats1. Biol. Reprod. 38, 1181–1188. 10.1095/biolreprod38.5.1181.

12. Anderson, E., and Albertini, D.F. (1976). Gap junctions between the oocyte and companion follicle cells in the mammalian ovary. J. cell Biol. 71, 680–686. 10.1083/jcb.71.2.680.

13. Makabe, S., Naguro, T., and Stallone, T. (2006). Oocyte–follicle cell interactions during ovarian follicle development, as seen by high resolution scanning and transmission electron microscopy in humans. Microsc. Res. Tech. 69, 436–449. 10.1002/jemt.20303.

14. El-Hayek, S., Yang, Q., Abbassi, L., FitzHarris, G., and Clarke, H.J. (2018). Mammalian Oocytes Locally Remodel Follicular Architecture to Provide the Foundation for Germline-Soma Communication. Curr Biol 28, 1124-1131.e3. 10.1016/j.cub.2018.02.039.

15. Eppig, J.J. (1976). Analysis of mouse oogenesis in vitro. Oocyte isolation and the utilization of exogenous energy sources by growing oocytes. J. Exp. Zoöl. 198, 375–381. 10.1002/jez.1401980311.

16. Eppig, J.J., Pendola, F.L., Wigglesworth, K., and Pendola, J.K. (2005). Mouse Oocytes Regulate Metabolic Cooperativity Between Granulosa Cells and Oocytes: Amino Acid Transport. Biol Reprod 73, 351–357. 10.1095/biolreprod.105.041798.

17. Su, Y.-Q., Sugiura, K., and Eppig, J. (2009). Mouse Oocyte Control of Granulosa Cell Development and Function: Paracrine Regulation of Cumulus Cell Metabolism. Semin Reprod Med 27, 032–042. 10.1055/s-0028-1108008.

18. Macaulay, A.D., Gilbert, I., Caballero, J., Barreto, R., Fournier, E., Tossou, P., Sirard, M.-A., Clarke, H.J., Khandjian, É.W., Richard, F.J., et al. (2014). The Gametic Synapse: RNA Transfer to the Bovine Oocyte1. Biol Reprod 91, 90. 10.1095/biolreprod.114.119867.

19. Albertini, D.F., Combelles, C.M., Benecchi, E., and Carabatsos, M.J. (2001). Cellular basis for paracrine regulation of ovarian follicle development. Reproduction Camb Engl 121, 647–653. 10.1530/rep.0.1210647.

20. Tang, S., Yang, N., Yu, M., Wang, S., Hu, X., Ni, H., and Cai, W. (2021). Autologous mitochondria transport via transzonal filopodia rejuvenates aged oocytes by UC-MSCs derived granulosa cells-oocyte aggregation. Biorxiv, 2021.10.30.466630. 10.1101/2021.10.30.466630.

21. Sha, H., Ye, Z., Ye, Z., Shi, S., Pan, J., Dong, X., and Zhao, Y. (2022). Germline-soma Supply Mitochondria for mtDNA Inheritance in Mouse Oogenesis. Biorxiv, 2022.01.11.475948. 10.1101/2022.01.11.475948.

22. Doherty, C.A., Tijjani, A., Munger, S.C., and Laird, D.J. (2025). Mammalian oocytes receive maternal-effect RNAs from granulosa cells. bioRxiv, 2025.02.10.637575. 10.1101/2025.02.10.637575.

23. Baena, V., and Terasaki, M. (2019). Three-dimensional organization of transzonal projections and other cytoplasmic extensions in the mouse ovarian follicle. Sci Rep-uk 9, 1262. 10.1038/s41598-018-37766-2.

24. Lewkowicz, E., Herit, F., Clainche, C.L., Bourdoncle, P., Perez, F., and Niedergang, F. (2008). The microtubule-binding protein CLIP-170 coordinates mDia1 and actin reorganization during CR3-mediated phagocytosis. J. Cell Biol. 183, 1287–1298. 10.1083/jcb.200807023.

25. Henty-Ridilla, J.L., Rankova, A., Eskin, J.A., Kenny, K., and Goode, B.L. (2016). Accelerated actin filament polymerization from microtubule plus ends. Science 352, 1004–1009. 10.1126/science.aaf1709.

26. Carabatsos, M.J., Elvin, J., Matzuk, M.M., and Albertini, D.F. (1998). Characterization of Oocyte and Follicle Development in Growth Differentiation Factor-9-Deficient Mice. Dev Biol 204, 373–384. 10.1006/dbio.1998.9087.

27. Zhang, Y., Wang, Y., Feng, X., Zhang, S., Xu, X., Li, L., Niu, S., Bo, Y., Wang, C., Li, Z., et al. (2021). Oocyte-derived microvilli control female fertility by optimizing ovarian follicle selection in mice. Nat Commun 12, 2523. 10.1038/s41467-021-22829-2.

28. Meng, W., Mushika, Y., Ichii, T., and Takeichi, M. (2008). Anchorage of microtubule minus ends to adherens junctions regulates epithelial cell-cell contacts. Cell 135, 948–959. 10.1016/j.cell.2008.09.040.

29. Tanaka, N., Meng, W., Nagae, S., and Takeichi, M. (2012). Nezha/CAMSAP3 and CAMSAP2 cooperate in epithelial-specific organization of noncentrosomal microtubules. P Natl Acad Sci Usa 109, 20029–20034. 10.1073/pnas.1218017109.

30. Jiang, K., Hua, S., Mohan, R., Grigoriev, I., Yau, K.W., Liu, Q., Katrukha, E.A., Altelaar, A.F.M., Heck, A.J.R., Hoogenraad, C.C., et al. (2014). Microtubule Minus-End Stabilization by Polymerization-Driven CAMSAP Deposition. Dev Cell 28, 295–309. 10.1016/j.devcel.2014.01.001.

31. Mora, J.M., Fenwick, M.A., Castle, L., Baithun, M., Ryder, T.A., Mobberley, M., Carzaniga, R., Franks, S., and Hardy, K. (2012). Characterization and Significance of Adhesion and Junction-Related Proteins in Mouse Ovarian Follicles. Biol Reprod 86, Article 153, 1–14. 10.1095/biolreprod.111.096156.

32. Gilula, N.B., Epstein, M.L., and Beers, W.H. (1978). Cell-to-cell communication and ovulation. A study of the cumulus-oocyte complex. J. cell Biol. 78, 58–75. 10.1083/jcb.78.1.58.

33. Duryee, W.R. (1954). SECTION OF BIOLOGY: MICRODISSECTION STUDIES ON HUMAN OVARIAN EGGS*. Trans. N. York Acad. Sci. 17, 103–108. 10.1111/j.2164-0947.1954.tb00399.x.

34. Rustom, A., Saffrich, R., Markovic, I., Walther, P., and Gerdes, H.-H. (2004). Nanotubular Highways for Intercellular Organelle Transport. Science 303, 1007–1010. 10.1126/science.1093133.

35. Clarke, H.J. (2022). Transzonal projections: Essential structures mediating intercellular communication in the mammalian ovarian follicle. Mol Reprod Dev. 10.1002/mrd.23645.

36. Zhang, C.-H., Liu, X.-Y., and Wang, J. (2023). Essential Role of Granulosa Cell Glucose and Lipid Metabolism on Oocytes and the Potential Metabolic Imbalance in Polycystic Ovary Syndrome. Int. J. Mol. Sci. 24, 16247. 10.3390/ijms242216247.

37. Matsunaga, H., Arikawa, K., Yamazaki, M., Wagatsuma, R., Ide, K., Samuel, A.Z., Takamochi, K., Suzuki, K., Hayashi, T., Hosokawa, M., et al. (2022). Reproducible and sensitive micro-tissue RNA sequencing from formalin-fixed paraffin-embedded tissues for spatial gene expression analysis. Sci. Rep. 12, 19511. 10.1038/s41598-022-23651-6.

38. Schindelin, J., Arganda-Carreras, I., Frise, E., Kaynig, V., Longair, M., Pietzsch, T., Preibisch, S., Rueden, C., Saalfeld, S., Schmid, B., et al. (2012). Fiji: an open-source platform for biological-image analysis. Nat. Methods 9, 676–682. 10.1038/nmeth.2019.

