## Supplementary figures for "Camsap3-Mediated Microtubules Maintain Transzonal Projections Essential for Germline-Soma Communication during Ovarian Follicle Maturation in Mice"

**Figure S1**

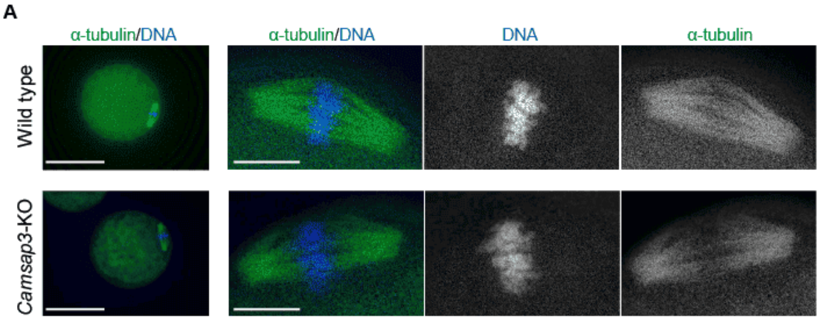


**Figure S1.**

(A) Oocytes obtained after superovulation were stained for α-tubulin and DNA. Although oocytes were rarely recovered from *Camsap3*-KO mice, the obtained oocytes appeared to form meiotic II spindles comparable to those in WT oocytes. Scale bars, 50 μm, 10 μm.

### Figure S2

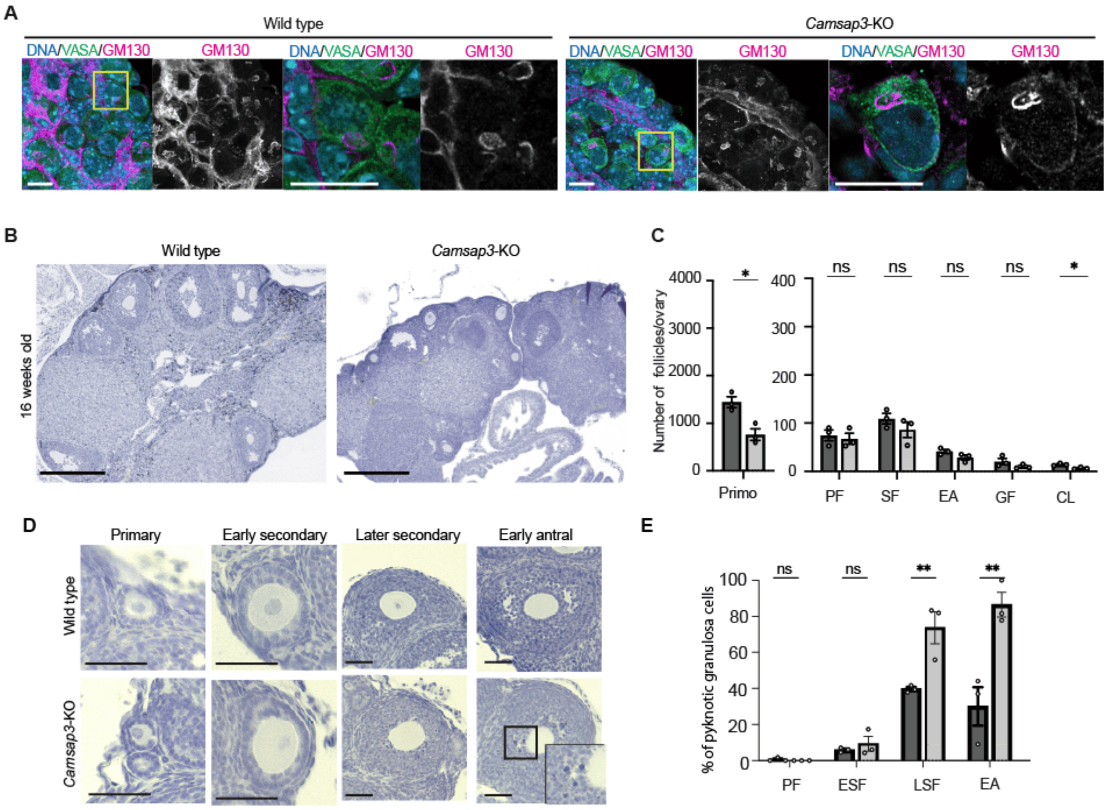


**Figure S2.**

(A) Representative images of densely packed primordial follicles from WT and *Camsap3*-KO mice. GM130 staining revealed a ring structure in oocytes of both WT and *Camsap3*-KO mice. Scale bar, 10 μm.

(B) Ovarian sections from 16-week-old WT and *Camsap3*-KO mice stained with haematoxylin. Scale bar, 300 μm.

(C) Average number of follicles at each stage per ovary: primordial (Primo), primary (PF), secondary (SF), early antral (EA), Graafian (GF) follicles and corpus luteum (CL) (16 weeks: WT, n=3; KO, n=3). *Camsap3*-KO mice had significantly fewer corpora lutea. Bars, mean; error bars, s.d.. *p < 0.05, two-tailed unpaired Student's t-test.

(D) Ovarian sections stained with haematoxylin showing pyknotic GCs (right inset). Scale bar, 50 μm.

(E) Proportion of follicles containing GCs with at least one pyknotic nucleus. (WT, n=3; KO, n=3) In late secondary and early antral follicles, *Camsap3*-KO mice showed an increased percentage of follicles undergoing regression. Bars, mean; error bars, s.d. *p < 0.05, **p < 0.01, two-tailed unpaired Student's t-test.

**Figure S3**

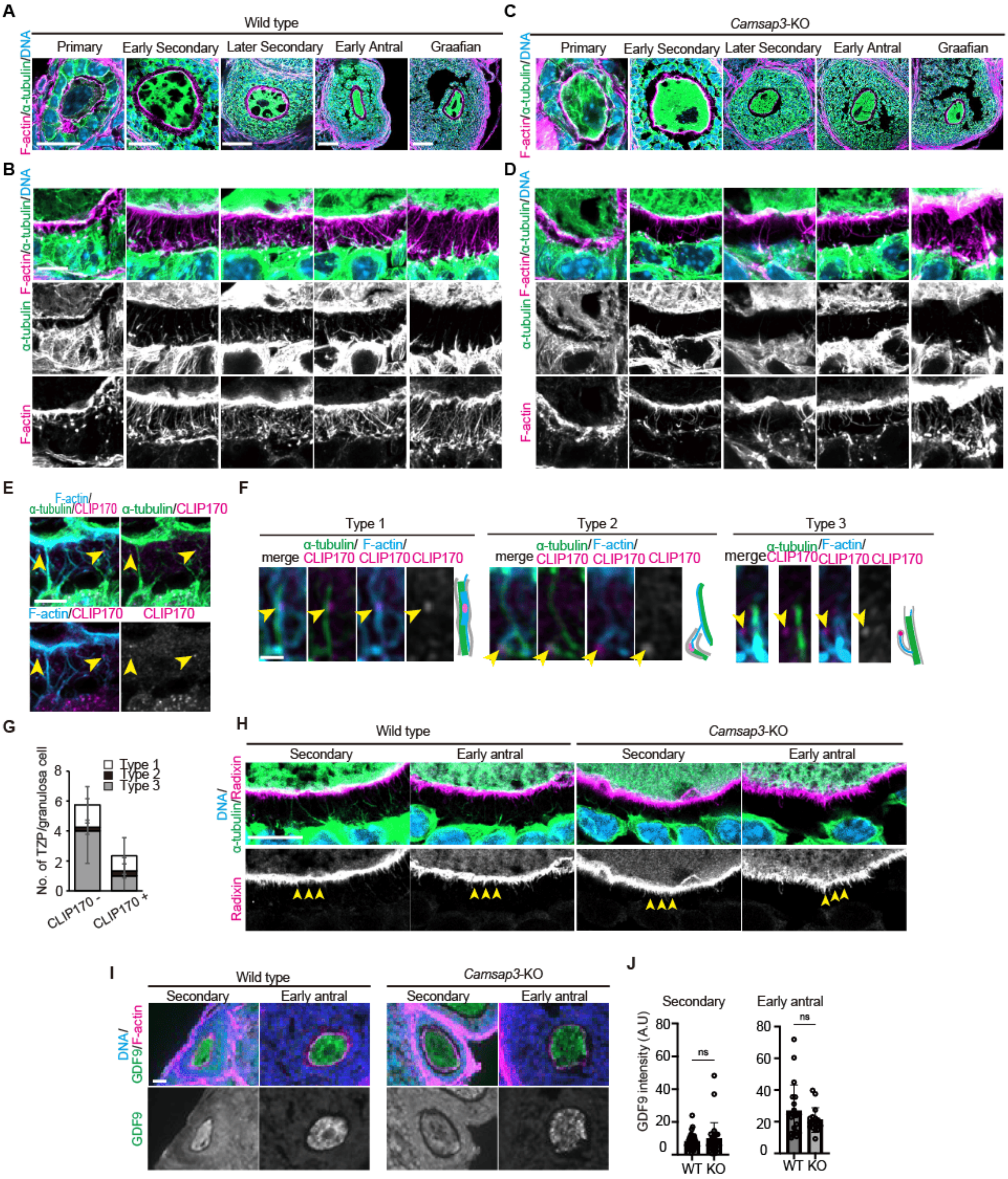


**Figure S3.**

(A, C) Representative images of follicles at each developmental stage. WT and *Camsap3*-KO ovarian sections were immunostained for F-actin, α-tubulin and DNA. Scale bar, 25 μm (primary, early secondary) 50 μm (later Secondary – Graafian).

(B, D) Magnified images of follicles at the primary, secondary, late secondary, early antral, and Graafian stages. Scale bar, 25 μm.

(E) Immunostaining for CLIP170, F-actin and α-tubulin in ovarian sections from WT mice. CLIP170 localised not only to microtubule tips but also to F-actin. Arrowheads indicate CLIP170 localisation to both microtubule and F-actin. Scale bar, 5 μm.

(F) CLIP170 localisation to TZPs according to the TZP-type defined in Figure 3. Scale bar, 0.5 μm.

(G) The mean number of TZPs per GC with or without CLIP170 localisation. TZPs with CLIP170 localisation were classified by type. Error bars, s.d.

(H) Immunostaining for radixin, α-tubulin and DNA in ovarian sections from WT and *Camsap3*-KO mice. Secondary and early antral follicles showed oocyte microvilli. In *Camsap3*-KO follicles, TZP numbers were reduced, whereas oocyte microvilli remained. Scale bar, 10 μm.

(I) Immunostaining for GDF9, F-actin and DNA in ovarian sections from WT and *Camsap3*-KO mice. GDF9 localised to the cytoplasm of the oocyte within the secondary and early antral follicles. Scale bar, 10 μm.

(J) The signal intensity of GDF9 in oocytes at the secondary and early antral stages. Bars, mean; error bars, s.d.

**Figure S4**

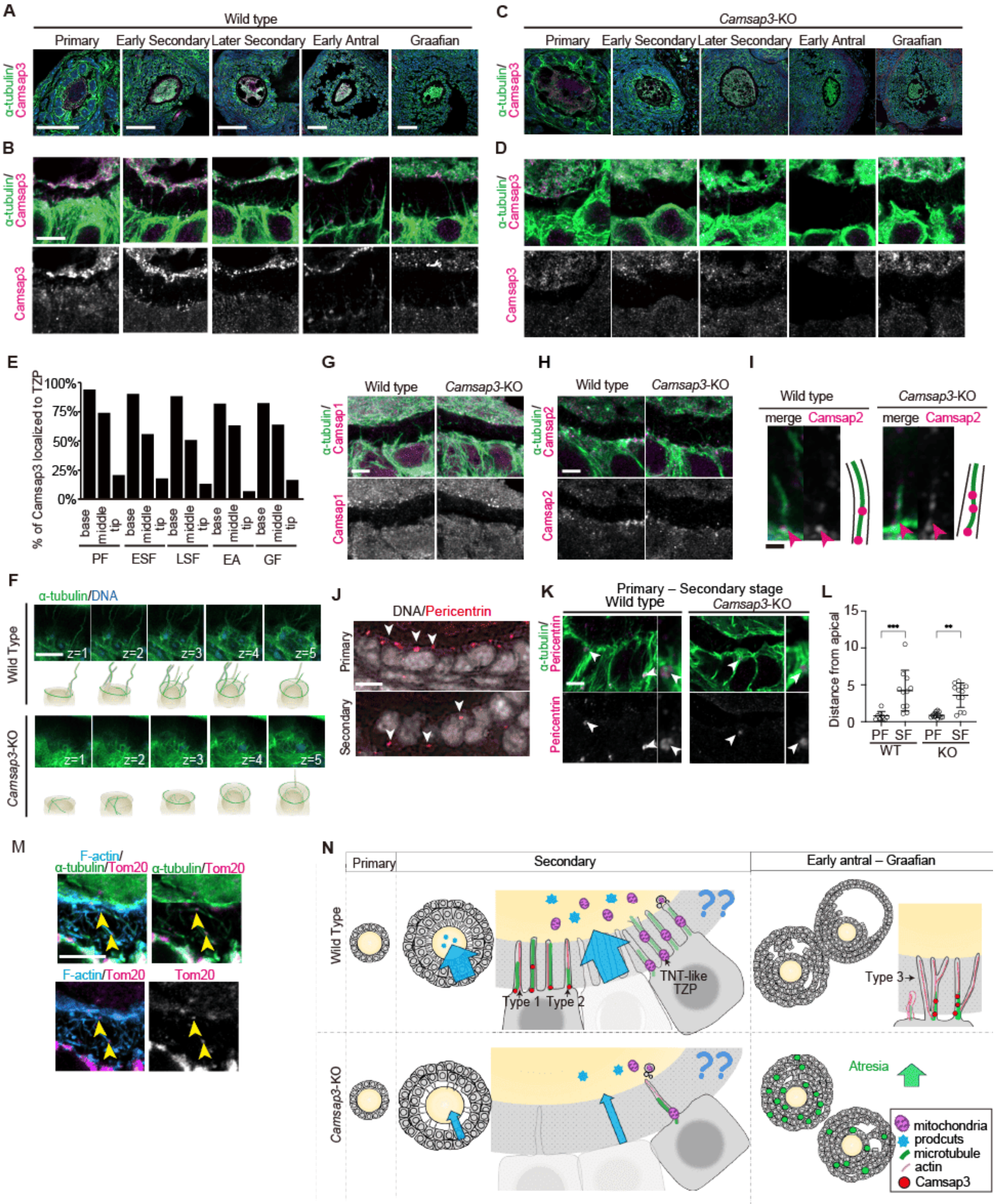


**Figure S4.**

(A, C) Immunostaining for Camsap3 and α-tubulin showing Camsap3 localisation to the apical surface of GCs in secondary and follicular follicles. Scale bars, 25 μm (primary) 50 μm (early secondary – Graafian) (B, D) Magnified images of the apical surface of GCs showing Camsap3 at the base and along TZPs (yellow arrows). Scale bar, 10 μm.

(E) Quantification of the three Camsap3 localisation patterns during follicle development.

(F) Apical views of GCs from isolated follicles. In WT, TZPs extended from the cell periphery towards the oocyte; The number of TZP was reduced in *Camsap3*-KO mice. Scale bar, 5 μm.

(G) No detectable Camsap1 was observed in GCs by immunostaining for Camsap1 and α-tubulin. Scale bar, 5 μm.

(H, I) Camsap2 localisation to GCs and TZPs indicated by immunostaining for Camsap2 and α-tubulin. Scale bar, 5 μm.

(J) Immunostaining for pericentrin and DNA revealed centrosomes at the apical surface in primary follicles, which were irregularly positioned within the granulosa layer in secondary and later stages. Scale bar, 5 μm.

(K) Representative images showing centrosomal microtubules encapsulated in TZPs. Scale bar, 0.5 μm.

(L) Distance of centrosome position from the apical surface of GCs.

(M) Immunostaining for Tomm20, F-actin and α-tubulin in WT ovaries. Arrowheads indicate mitochondria detected along TZP microtubules. Scale bar, 5 μm.

(N) Model of Camsap3 function in ovarian follicles: microtubules play a more important role than previously thought in TZP maintenance, which is essential for follicle development. Camsap3-mediated microtubules support the temporal transition of TZP morphology during follicle development. In the absence of Camsap3, the number of TZPs was significantly reduced, including TNT-like TZPs, which may facilitate the translocation of macromolecules and organelles, such as mitochondria.
